# Postnatal development of pyramidal neurons excitability and synaptic inputs in mouse gustatory cortical circuits

**DOI:** 10.1101/2025.09.11.675590

**Authors:** Hillary Schiff, Arianna Maffei

## Abstract

Cortical neurons in sensory areas undergo a protracted process of postnatal maturation that includes changes in membrane properties, synaptic drive and connectivity. The completion of this process is associated with the closure of sensitive windows for experience-dependent plasticity. Weaning is a critical time in the development of taste circuits as animals transition from depending on the mother for nutrition to eating independently. While there is some evidence for developmental changes in taste bud innervation and in membrane properties of neurons in brainstem circuits for taste, very little is known about postnatal changes in the gustatory cortex (GC), the primary cortical region for taste and taste-guided behaviors. Here, we focused on pyramidal neurons in the deep layers of GC and compared their membrane properties in pre- and postweaning age groups. We report dynamic changes in intrinsic excitability and a progressive shift of the excitation/inhibition (E/I) balance toward inhibition as pyramidal neurons reach their young adult properties. The increase in inhibitory drive accompanied a protracted process of postnatal maturation of inhibitory circuits mediated by parvalbumin expressing neurons (PV neurons) that showed a progressive increase in their association with perineuronal nets (PNNs) and refinement of their connectivity onto pyramidal neurons. Together, our results indicate that GC neurons undergo protracted postnatal maturation that may influence taste response properties at the transition to independent feeding.

**Significance Statement:** We show that the circuit in the gustatory cortex undergoes a protracted maturation process that extends into adulthood and progressively shifts the excitability of GC toward inhibition through changes in pyramidal neurons membrane properties, increased inhibitory synaptic drive and refinement of parvalbumin neurons connectivity. This maturation process extends beyond the developmental windows previously reported for other sensory cortical circuits and overlaps with the recently identified critical period for the development of taste preferences. As finding nutritious food sources may require the integration of vision, audition, somatosensation and olfaction, the extended maturation process for taste cortical circuits may facilitate the integration of sensory information for the identification of food and the decision to ingest it.

## Introduction

Weaning is an important transition in mammalian development, as animals shift from relying on their mother’s milk to foraging for food. This early experience with feeding independence influences the development of taste preferences (Schiff et al., 2023). While the postnatal development of gustatory cortical circuits is not well studied, there is some experimental evidence for protracted maturation of neuronal morphology and early-life experience-dependent effects on neurons in other regions of the taste system. For instance, postnatal anatomical re-wiring has been observed in the first central relay in the gustatory system, the nucleus of the solitary tract (NTS). This process occurs after weaning, with the inputs to the NTS reaching mature connectivity by postnatal day 35 (P35) and undergoing additional refinement into adulthood (Hill et al., 1983; Sollars et al., 2006; May et al., 2008; Sun et al., 2017). In the gustatory portion of the parabrachial nucleus, dendritic arborization of multipolar and fusiform cells reach adult morphology by P35 (Lasiter and Kachele, 1988).

In primary sensory cortices, including vision, audition, and somatosensation, developmental time windows of heightened plasticity have been identified as sensitive periods (Micheva and Beaulieu, 1995; Maffei et al., 2006; Maffei and Turrigiano, 2008; Maffei et al., 2010; Wang et al., 2011; Takesian et al., 2012; Gainey and Feldman, 2017; Gainey et al., 2018; Takesian et al., 2018). During these periods, cortical circuits undergo a series of maturation processes and show heightened sensitivity to sensory inputs.

GABAergic inhibitory synapses in particular play a crucial role in postnatal circuit maturation and refinement. Inhibitory circuits themselves undergo protracted postnatal maturation (Hensch, 2004; Tatti et al., 2017; Takesian et al., 2018), with increases in GABAergic inhibition opening the critical period for circuit refinement. For instance, in a knock-out mouse in which GABA is severely diminished (GAD-KO), the critical period may never open unless GABA receptors are activated pharmacologically (Fagiolini and Hensch, 2000). Experience-dependent maturation of inhibitory circuits during sensitive developmental windows is primarily associated with changes in the parvalbumin-expressing subtype of interneuron (PV^+^ INs). Over the course of postnatal development, there is an increase in the number of PV^+^ INs (Gonchar et al., 2007; Tatti et al., 2017) along with perisomatic innervation of pyramidal neurons by fast-spiking basket neurons (Chattopadhyaya et al., 2004). This process is associated with increases in the expression of PV in PV^+^ INs (Murase et al., 2017) and with the accumulation of perineuronal nets (PNNs) around PV^+^ INs somata. The presence of mature PNNs correlates with the duration of the critical period (Sorg et al., 2016).

In adult GC, inhibitory responses make up a component of the complex taste-evoked responses (Yamamoto, 1984) and shape the excitability of pyramidal neurons. Furthermore, feedforward inhibition mediated by PV^+^ INs is recruited by both thalamocortical (Haley et al., 2023) and basolateral amygdala inputs (Haley et al., 2016), suggesting that this population of inhibitory neurons shapes the response properties of GC pyramidal neurons. Recent work identified a sensitive window for taste experience-dependent plasticity that depends on the maturation of PV^+^ INs (Schiff et al., 2023). This suggests that despite significant differences in circuit organization (Haley et al., 2016, 2023), GC may share some mechanisms of postnatal maturation with other cortical regions.

In this study, we used morphological reconstructions, whole-cell patch-clamp electrophysiology in acute slice preparations, immunohistochemistry, and channelrhodopsin assisted circuit mapping (CRACM (Petreanu et al., 2007)) to track postnatal changes in the membrane properties of deep layer pyramidal neurons and assess possible age-dependent refinement of the connectivity of GC PV^+^ INs of male and female mice during the postnatal window straddling the transition to independent feeding and adulthood. Pyramidal neurons showed a transient, post-weaning shift in input/output function and a progressive shift in the excitation/inhibition (E/I) balance toward inhibition mediated due to an increase in inhibitory synaptic drive with no overall change in excitation. The changes in synaptic inhibition were accompanied by increases in PV fluorescence intensity, accumulation of PNNs around PV^+^ INs and refinement of the connectivity of PV^+^ INs around the perisomatic region of pyramidal neurons. Our results point to a dynamic circuit refinement process occurring at the transition to independent feeding and extending into adulthood, a period that overlaps with the sensitive window for the development of sweet taste preference.

## Materials and Methods

### Animals

All experimental procedures followed the guidelines of the National Institute of Health and were approved by the Institutional Animal Care and Use Committee. Mice of both sexes were group-housed in a vivarium on a 12-hour light-dark cycle. Experiments were performed during the light cycle. Wild-type C57BL/6 mice were purchased from Charles River, arriving to our facility either as adults, or as litters, consisting of the lactating mothers and pups at postnatal day 5 (P5). PV-cre (JAX stock #017320; (Taniguchi et al., 2011)) female and Ai14 (RRID: IMSR_JAX:007908) male mice, both homozygous, were ordered from Jackson Laboratory and crossed in our animal facility. The Ai14 line expresses tdTomato in a cre-dependent fashion (Madisen et al., 2010). The PV-cre;Ai14 offspring were heterozygous for both cre and lox-stop-lox-tdTomato alleles and were used in CRACM experiments.

### Stereotaxic Surgery

Mice aged P14 ± 1 or P49 ± 1 were deeply anesthetized with a cocktail of 100mg/kg ketamine and 10mg/kg xylazine injected IP. Once anesthetized, they were placed on the stereotaxic apparatus and received an injection of bupivacaine (2.5mg/ml, ∼0.1ml) under the scalp for local anesthesia. A craniotomy was made over the left GC at coordinates adjusted for developmental stage (P14: +0.9mm anterior to Bregma, +3.05mm lateral from Bregma; P49: +1.0mm anterior to Bregma, +3.25mm lateral from Bregma). A pulled glass micropipette filled with mineral oil was attached to a Nanoject II (Drummond) and backfilled with a solution containing a viral construct (AAV9-hSyn-FLEX-soCoChR). To reach the appropriate titer of viral particles, the stock virus was diluted 1:5 in sterile saline. The tip of the glass micropipette was slowly lowered to a depth of 2.3mm below the pial surface for both age groups. Approximately 250nl of the virus solution was injected with pressure pulses delivering approximately 13.8nl per pulse over approximately 10min. To ensure localized delivery of the viral construct, the pipette tip was left in place for approximately 10min after the end of the injection, then slowly retracted. The skin over the craniotomy was then sutured closed and covered with vetbond. Mice recovered on a heating pad until ambulatory at which point they were returned to the home cage.

### Electrophysiology

To prepare acute coronal slices (350µm) containing GC mice were anesthetized with isoflurane using the bell jar method and rapidly decapitated. The brain was dissected in ice cold, oxygenated standard artificial cerebrospinal fluid (ACSF) containing, in mM: 126 NaCl, 3 KCl, 25 NaHCO_3_, 1 NaHPO_4_, 2 MgSO_4_, 2 CaCl_2_, 14 dextrose with an osmolarity of 313-317mOsm with a pH = 7.4 when bubbled with carbogen (95% oxygen, 5% carbon dioxide). The brains were secured to a fresh tissue vibratome (Leica VT1000) and were submerged in oxygenated ACSF during the slicing procedure. Slices were recovered in 34°C oxygenated ACSF for 20min and then brought to room temperature for 30min before beginning recordings. Individual slices were transferred to the recording chamber mounted on an upright microscope (Olympus BX51WI). To record input/output functions, slices were continuously perfused with oxygenated ACSF and maintained at 34°C with an inline heater (Harvard Bioscience). Whole-cell patch clamp recordings in current clamp were obtained from visually identified pyramidal neurons under DIC optics using borosilicate glass pipettes with resistance of 3–5MΩ.

The internal solution contained, in mM: 100 K-Gluconate, 20 KCl, 10 K-HEPES, 4 Mg-ATP, 0.3 Na-GTP, 10 Na-phosphocreatine, and 0.4% biocytin, pH was adjusted to 7.35 with KOH; osmolarity was adjusted to 295mOsm with sucrose. Neurons with a series resistance >15MΩ or those with series resistance changing >20% during recording were excluded from the analysis. Neuron identity was confirmed *post hoc* with immunohistochemistry aimed at reconstructing neuron morphology, verifying laminar location, and assessing lack of expression of the GABA neuron marker GAD67. Current steps (700ms) of increasing intensity (-100 to 400pA at 50pA increments) were injected into the cell with a 10s inter–sweep interval. The dynamic input resistance (DIR) was calculated as the slope of the current-voltage curve for steps below action potential threshold. Input-output curves represent the frequency of action potential firing for each suprathreshold current step averaged across recorded neurons for each age group. Rheobase and AP threshold were quantified from a trace in which the neuron fired a single action potential for 2 consecutive sweeps. This was accomplished by injecting current steps (700ms) with increasing amplitude (2pA steps) around the suprathreshold range. The AP threshold was measured as the voltage at which the first derivative of the voltage trace reaches 20mV/ms. For a small number of neurons, it was not possible to isolate 2 consecutive traces with a single AP, preventing reliable rheobase and AP threshold measurements. This is reflected in the reported number of neurons used for statistical comparisons in the Results section.

To record spontaneous synaptic currents, slices were perfused with an oxygenated ACSF solution optimized to facilitate spontaneous activity (38), in mM: 124 NaCl, 3.5 KCl, 26 NaHCO3, 1.25 NaHPO4, 0.5 MgCl2, 1 CaCl2, 14 dextrose, maintained at 34°C with an inline heater (Harvard Bioscience). Recordings were obtained in voltage clamp with an internal solution containing in mM: 20 KCl, 100 Cs-sulfate, 10 K-HEPES, 4 Mg-ATP, 0.3 Na-GTP, 10 Na-phosphocreatine, 0.2% biocytin (Vrev [Cl^−^] = −49.3 mV). The pH was adjusted to 7.35 with KOH, and the osmolarity was adjusted to 295mOsm with sucrose. The sodium channel blocker QX314 (3mM, Tocris Bioscience) was added to the internal solution to stabilize recordings during prolonged depolarization. Spontaneous postsynaptic excitatory and inhibitory currents (sEPSCs and sIPSCs) were recorded by holding neurons at three different holding potentials around the expected reversal potential for chloride (in mV: −55, –50, −45) and for AMPA/NMDA receptor mediated currents (in mV: +5, +10, +15). *Post hoc* current vs. holding voltage (Vhold) functions were used to identify the holding voltage that best isolated sEPSCs and sIPSCs respectively. Traces recorded at the optimal holding voltage were then used for analysis of amplitude and frequency of spontaneous events (12, 38). Individual postsynaptic events were identified using template search method in Clampfit (Clements and Bekkers, 1997). Cumulative distributions for sEPSCs amplitude and frequency included 35 events/cell, while cumulative distributions for sIPSCs included 100 events/cell. The difference in number of events included in the cumulative distributions was due to the overall lower frequency of sEPSCs compared to sIPSCs in infragranular GC pyramidal neurons. These events were then averaged and rise time and decay time constants were calculated for each cell. Total charge was calculated in IGOR Pro as the area under the curve for a recording period of 5 minutes.

### Channelrhodopsin assisted circuit mapping

A virus coding for a cre-dependent soma-restricted channelrhodopsin (AAV9-hSyn-FLEX-soCoChR-GFP, titer ∼1e^13^ vg/mL Addgene cat # 107712-AAV9) was injected into the left GC of PV-cre;Ai14 mice and allowed to incubate for 1 week after verification that this was the optimal time to allow for sufficient expression of the opsin while maintaining soma-restriction. Red fluorescence in PV^+^ neurons allowed us to direct our recordings to an area where PV^+^ neurons were present. Initial tests informed us that expression was very low in P17 mice following virus injection at P10, consistent with the small number of PV^+^ INs quantified at this age with immunohistochemistry (see Fig. 4). We reasoned that this occurred because PV expression and the number of PV^+^ neurons was too low for sufficient cre expression and thus cre-dependent recombination and expression of the viral products. Thus, to assess connectivity from PV^+^ INs onto pyramidal neurons around weaning time and compare it to that of adult mice, we performed virus injection at P14 ± 1 and recordings at P21 ± 1. For the older age group, stereotaxic surgery for virus injection was at P49 ± 1 and recordings at P56 ± 1.

We recorded from visually identified neurons that did not display red fluorescence and were putatively PV^-^. *Post hoc* biocytin staining confirmed that these neurons were negative for tdTomato and their location and morphology was used to confirm that they were indeed infragranular pyramidal neurons. We delivered patterned light stimulation using a digital micromirror device (Polygon DMD device, Mightex) to focus light from a blue LED (473nm) onto ∼100×100µm square regions of interest (ROIs) through a 4x objective (Olympus). Pulses were delivered at 1Hz in an ordered sequence of 5×7 ROIs spanning the cortical column surrounding the recorded neuron (approximately 500µm dorsal to ventral and 700µm lateral to medial beginning at the border of layers 1 and 2 and extending to the white matter/corpus collosum). Pulse duration was 5ms. Light intensity was determined by initial stimulation of the full field of view using an intensity that elicited a reliable postsynaptic response. We collected 10 sequences of 5×7 ROI stimulation and averaged the traces in each ROI to determine the magnitude of the IPSC evoked in each illumination region. Although 35 ROIs were stimulated, analyses were conducted on only 30 ROIs because the last 5 ROIs (closest to the white matter) often stimulated outside of GC into the claustrum or within the white matter. As none of our recorded neurons responded to stimulation of these 5 ROIs, they were excluded from analysis. Neurons were considered responsive to stimulation of a specific ROI if an IPSC was elicited in at least 3 out of the 10 sweeps.

### Labeling of recorded neurons

Following electrophysiological recordings, slices were fixed in 4% PFA for 24 hours to 1 week. They were then washed in PBS at RT (3 × 5min) and blocked in 5% NGS and 5% BSA in PBST (1.0% Triton X-100) for 2-4h at RT. Next, slices were incubated with primary antibodies overnight at 4°C in a solution containing 1% NGS and 1% BSA in PBST (0.1% Triton X-100). The following antibodies were used: streptavidin Alexa Fluor-568 conjugate or streptavidin Alexa Fluor-647 conjugate (1:2000, Invitrogen, S11226 or S32357, respectively), mouse anti-GAD67 (1:500, MilliporeSigma, MAB5406, monoclonal), and/or rabbit anti-RFP (1:500, abcam 390 004). Sections were then washed with PBS (3 × 10min) and incubated with at RT for 4h with goat anti-mouse Alexa Fluor 488 (1:200, Thermo Fisher A-11029) and/or goat anti-rabbit Alexa Fluor 568 (1:200 Thermo Fisher A-11011), and counterstained with Hoechst 33342 (1:5000, Invitrogen, H3570). After washing with PBS (3 × 10min), sections were mounted onto glass slides with Fluoromount-G. Sections were imaged with a laser-scanning confocal microscope (Olympus Fluoview) at 10x magnification for validation of location in infragranular GC and at 40x to determine absence of colocalization with GAD67.

A subset of recorded neurons was first imaged with fluorescent labeling as described above and then converted to 3,3’-Diaminobenzidine (DAB) for anatomical tracing with Neurolucida. To do this, coverslips were removed from the slides by soaking in PBS and the tissue was removed from the slides. The tissue was rinsed thoroughly in PBS at RT (6 x 10min). Slices were incubated in 0.3% H_2_O_2_ at 4°C for 1 hour and then rinsed in PBS (4 x 10min). An Avidin-Biotin complex (Vectastain Elite ABC HRP Kit-Vector Labs PK-6100) was kept in the dark for 30min and then brought up to 10mL total volume with PBS and 0.1% Triton X-100. Slices incubated in this solution in the dark at 4℃ overnight. The following day, after 4 x 10min PBS rinses, slices were pre-incubated in a DAB solution (Vectastain DAB Peroxidase (HRP) Substrate Kit (with nickel), 3,3’-diaminobenzidine-Vector Labs SK-4100) at room temperature for 15min, before adding H_2_O_2_ to the well with constant shaking for 40-60s, until the brown signal was visible by eye. The tissue was quickly rinsed with PBS followed by 4 x 10min rinses with phosphate buffer (PB). Once mounted on the slides, slices were allowed to dry and then dehydrated through a series of increasing concentrations of ethanol followed by Xylene and coverslipped with Entallan medium.

### Immunohistochemistry for PV and PNN labeling

Mice were deeply anesthetized and transcardially perfused with phosphate-buffered saline (PBS), followed by perfusion with 4% paraformaldehyde (PFA) in PBS. Brains were dissected and postfixed in 4% PFA at 4°C for a minimum of 3h to a maximum of 24h followed by incubation in PBS-buffered sucrose (30%) solution for cryoprotection until brains were saturated (∼36h). Thin (50µm) coronal sections containing GC were cut on a vibratome (VT1000, Leica). Brain sections were first washed in PBS at RT (3 × 5min), then incubated in blocking buffer (5% normal goat serum (NGS) and 5% bovine serum albumin (BSA) in PBST with 0.5% Triton X-100) for 1h at RT. Next, sections were incubated with primary antibodies (mouse anti-PV, 1:2000, Swant 235, RRID: AB_10000343; biotinylated wisteria floribunda lectin, WFA, 1:500, Vector Laboratories B-1355-2) overnight at 4°C in a solution containing 2% NGS and 1% BSA in PBST (0.1% Triton X-100). Sections were then washed with PBS (3 × 10min) and incubated with the following fluorescent secondary antibodies at RT for 2h: goat anti-mouse Alexa Fluor 488 (1:200, Thermo Fisher A-11029), streptavidin Alexa Fluor 647 (1:500, Thermo Fisher S32357) and counterstained with neuronal-targeting fluorescent Nissl (Neurotrace530/615, 1:200 Thermo Fisher N21482). After washing with PBS (3 × 10min), sections were mounted onto glass slides with Fluoromount-G (Southern Biotech). Images were obtained with a laser-scanning confocal microscope (Olympus Fluoview) at 10x magnification.

### PV and WFA imaging analysis

For each animal, we quantified PV and PNN signals in two sections containing GC spaced at 150µm. The GC outline was previously created and aligned for each section using only the neurotrace counterstain to avoid potential bias that might emerge when observing the PV and WFA signal. Quantification of the number of PV^+^ INs and PNNs as well as of fluorescence intensity was obtained in ImageJ using the Pipsqueak AI macro from Rewire Neuro, Inc (Sorg et al., 2016). Individual ROIs for each PV^+^ neuron or PNN were automatically created and verified by an experimenter who was blind to the experimental condition. All single- or double-labeled neurons or PNNs were detected and analyzed. The proportion of PV^+^ INs was quantified relative to the fluorescent Nissl counterstain (Neurotrace). Quantification of neurotrace-positive cells was performed using the ImageJ plugin image-based tool for counting nuclei (ITCN; Center for Bio-image Informatics, University of California, Santa Barbara).

### Digital morphological reconstruction

Digital reconstruction of recorded pyramidal neurons was performed in Neurolucida (MBF Bioscience) at 40x magnification. Neurons were deemed suitable for reconstruction if they had at least a tuft of basal dendrites. The soma and all visible basal and apical dendrites were traced. Neurolucida Explorer was used for analysis of each cell for the following parameters: number of basal dendrites, total basal dendrite length, number of basal nodes, total apical dendrite length, number of apical nodes, and convex hull.

### Data analysis

Data are presented as mean ± standard error of the mean (SEM). Electrophysiological analyses for intrinsic properties and opsin-elicited postsynaptic responses were performed in Igor Pro (WaveMetrics). Cumulative distributions for sEPSCs and sIPSCs used the event detection template search in Clampfit (Molecular Devices). Statistical comparisons were made in Graphpad Prism version 10. Statistical tests applied to each data set are specified in the Results. Statistical significance was determined as a p-value ≤ 0.05.

## Results

We quantified intrinsic excitability, neuronal morphology and synaptic drive onto infragranular pyramidal neurons recorded from the gustatory cortex (GC). Pyramidal neurons were visually identified under DIC optics. Upon achieving the whole cell configuration, square current steps (700ms) of increasing amplitude were injected to assess the electrophysiological signature of the recorded neuron and obtain the input/output function in current clamp. A group of pyramidal neurons were recorded in voltage clamp while holding neurons at the expected reversal potential for excitatory and inhibitory currents to quantify synaptic drive. All recorded neurons were filled with biocytin for *post hoc* morphology reconstructions using fluorescence microscopy. Neurolucida tracing was used to reconstruct a subset of recorded neurons for detailed analysis of neuronal morphology. Channelrhodopsin assisted circuit mapping (CRACM, (Petreanu et al., 2007)) was used to determine postnatal developmental changes in the recurrent connectivity of perisomatic inhibition mediated by PV^+^ neurons.

### Changes in GC pyramidal neurons excitability and morphology during postnatal development

We began the analysis of pyramidal neurons membrane properties by quantifying their intrinsic excitability in the presence of blockers for ionotropic glutamate and GABA receptors. Patch clamp recordings in the current clamp configurations were obtained from pyramidal neurons in acute coronal slices containing GC prepared from C57BL/6 mice at P17, P35, and P56 to straddle the postnatal developmental window from pre-weaning to young adulthood. All recorded neurons were confirmed GC pyramidal neurons in the infragranular layers as indicated by *post hoc* fluorescence histological procedures (**Fig. 1A**) and their firing pattern (**Fig. 1B**). Recordings were from all three divisions of GC (granular, dysgranular and agranular). As the analysis did not show differences in membrane properties across divisions, data were pooled. Input/output functions were built by plotting the frequency of action potentials in response to depolarizing current steps of increasing amplitude (50pA increments) for neurons recorded in slices from each age group. We observed a transient flattening of the input-output curve between P17 and P35. This effect was reversed in neurons recorded at P56, which showed an input/output function comparable to that of P17 mice (**Fig. 1C**, input-output: P17 n=22 cells from 8 mice; P35 n=22 cells from 11 mice; P56 n=11 cells from 5 mice; F_(2,52)_=3.85 p=0.028, 2-way repeated measures (RM) ANOVA with main effect of age). There was also a significant but transient reduction in the maximum action potential firing frequency from P17 to P35, followed by a recovery to P17 levels at P56 (**Fig. 1D**, max AP firing: P17 n=22 cells from 8 mice; P35 n=22 cells from 11 mice; P56 n=11 cells from 5 mice; F_(2,52)_=5.33 p=0.01, one-way ANOVA with Tukey’s *post hoc* test: P17 vs P35 p=0.01, P17 vs P56 p=0.99, P35 vs P56 p=0.05). These effects were not accompanied by changes in action potential threshold, nor rheobase, consistent with a lack of a lateral shift in the input/output function (**Fig. 1E**, AP threshold: P17 n=20 cells from 8 mice; P35 n=18 cells from 10 mice; P56 n=8 from 4 mice; F_(2,43)_=2.843 p=0.0693; **Fig. 1F**, rheobase: P17 n=22 cells from 8 mice; P35 n=18 cells from 10 mice; P56 n=8 from 4 mice; F_(2,45)_=1.761 p=0.4624, one-way ANOVA).

**Figure 1.**
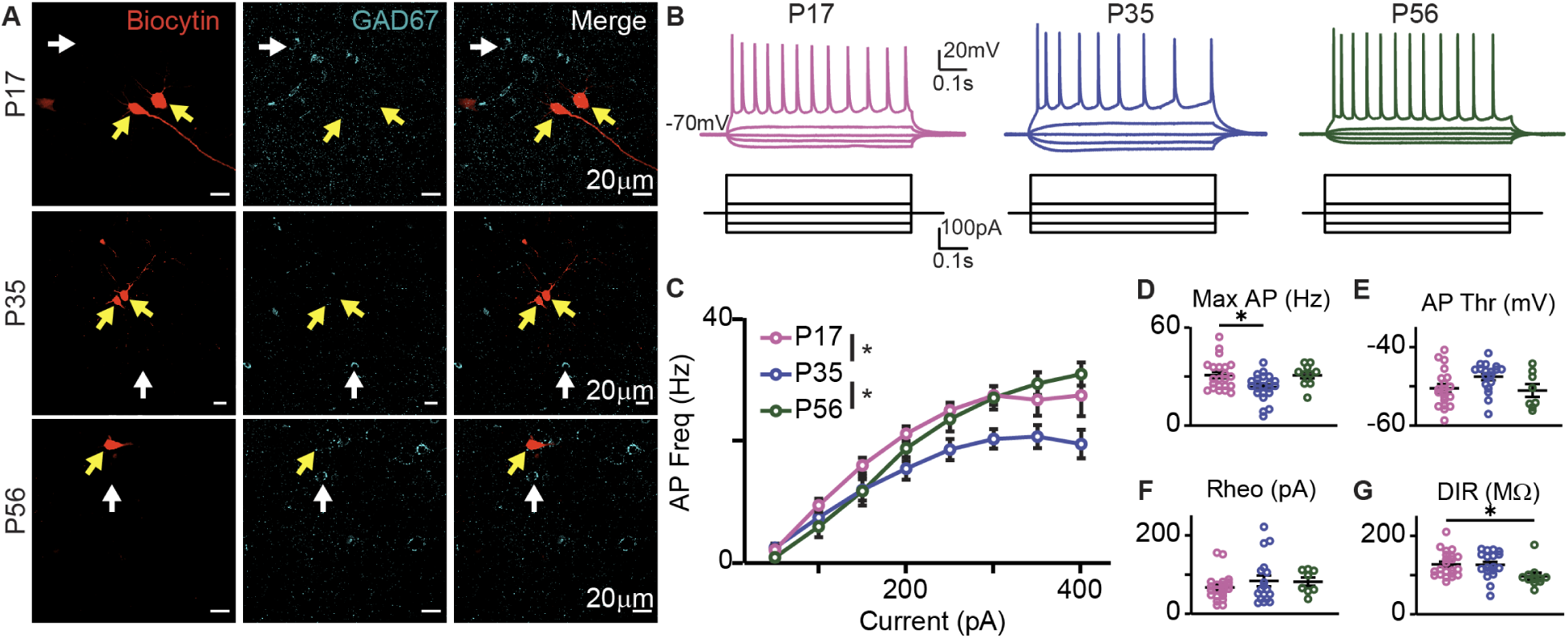
Postnatal developmental changes in intrinsic excitability of GC infragranular pyramidal neurons. **A.** Sample single confocal plane images of biocytin-filled recorded neurons showing lack of colocalization with GAD67. The rows show a sample recorded neuron for each age. The yellow arrows point to the recorded neuron filled with biocytin (red); the white arrows point to examples of GAD67^+^ neurons at the same tissue depth. The lack of overlap indicates that the recorded neuron is GAD67^-^. **B.** Top. Sample traces of pyramidal neurons recorded at P17 (pink), P35 (blue) and P56 (green). Bottom. Diagram of step current injections. **C.** Average input/output function for neurons recorded at each age. Data presented as mean ± SEM. **D.** Maximum firing frequency for all recorded neurons plotted by age group. **E.** Action potential (AP) threshold for each recorded neuron by age group. **F.** Rheobase for each recorded neuron in each age group. **G.** Dynamic input resistance (DIR) quantified as the slope of the input/output function for steps below rheobase. For D-G: black horizontal lines: average values ± SEM; open circles: single neuron measurements. Asterisks: p ≤ 0.05.

Interestingly, despite the transient shift in input/output function, the dynamic input resistance (“DIR”) showed a small but significant and progressive decrease from pre-weaning age (P17) to adulthood (**Fig. 1G**, DIR; P17 n=22 cells from 8 mice; P35 n=18 cells from 8 mice; P56 n=11 cells from 5 mice; F_(2,48)_=3.721 p=0.032, one-way ANOVA with Tukey’s *post hoc* test: P17 vs P35 p=0.98, P17 vs P56 p=0.04, P35 vs P56 p=0.05). Overall, these results suggest that the intrinsic excitability of GC infragranular pyramidal neurons decreases during the 5th postnatal week. This effect is transient, as the input/output function and maximum firing frequency return to pre-weaning levels by early adulthood, despite a significant drop in dynamic input resistance.

Changes in dynamic input resistance may depend on intrinsic membrane properties or on the complexity of neuronal morphology, which tends to increase during postnatal development (Elston and Fujita, 2014; Ciganok-Huckels et al., 2023). To quantify dendritic morphology, we converted a subset of fluorescently labeled neurons included in the analysis of intrinsic properties to DAB staining. The pyramidal morphology and deep layer location of the cell body had already been verified with fluorescent imaging overlayed with Neurotrace to identify cortical layers. Slices reprocessed for DAB staining facilitated the visualization of processes across visual planes and their reconstruction with Neurolucida software while avoiding photobleaching (**Fig. 2A**). For each neuron, we quantified the number of dendritic branches, dendritic length, volume of their dendritic arborizations, and dendritic complexity (**Fig. 2B**). It should be noted that all neurons included in this morphological analysis showed comparable electrophysiological signature (see **Fig. 1**) and apical dendrites extending tufted arborizations into layer 1, consistent with the typical morphology of infragranular pyramidal neurons. The average distance of cell bodies from the pial surface and the average length of apical, but not basal, dendrites were significantly increased in GC neurons in P56 mice compared to P17 and P35 suggesting an increase in cortical thickness from P17 to P56 (**Fig. 2C**, somata distance from the pia: P17 n=5 neurons; P35 n=15 neurons; P56 n=7 neurons; F_(2,24)_=6.892 p=0.0043 one-way ANOVA with Tukey’s *post hoc* test: P17 vs P35 p=0.3481; P17 vs P56 p=0.0049; P35 vs P56 p=0.0211; **Fig. 2D**, apical dendrite length: P17 n=3 neurons; P35 n=5 neurons; P56 n=3 neurons; F_(2,8)_=20.03 p=0.0008; *post hoc*: P17 vs P35 p=0.9437; P17 vs P56 p=0.0016; P35 vs P56 p=0.0011; **Fig. 2E**, basal dendrites length: P17 n=4 neurons; P35 n=14 neurons; P56 n=7 neurons; F_(2,22)_=0.5585). Consistent with this possibility, convex hull analysis revealed a greater volume of arborizations at P56 compared to younger ages (**Fig. 2F**, convex hull arborization: P17 n=2 neurons; P35 n=5 neurons; P56 n=3 neurons; F_(2,7)_=5.832 p=0.0323 one-way ANOVA with Tukey’s *post hoc* test: P17 vs P35 p=0.9773; P17 vs P56 p=0.0689; P35 vs P56 p=0.0377). Differently, the total number of dendrites and the dendritic complexity, quantified with Sholl analysis, did not change (**Fig. 2G-H**, number of dendrites: P17 n=4 neurons; P35 n=14 neurons; P56 n=7 neurons; F_(2,22)_=1.061 p=0.3630, one-way ANOVA; Sholl analysis: P17 n=4 neurons; P35 n=14 neurons; P56 n=7 neurons; F_(2,22)_=0.4696 p=0.9474, two-way RM ANOVA with main effect of age). These data suggest that despite the increase in length of the apical dendrite and the increased volume, the complexity of pyramidal neuron morphology is largely stable across the ages we examined. Together, the analysis of intrinsic properties and morphology suggests that the transient changes in input/output function are not explained by substantial changes in neuronal morphology and are likely due to postnatal maturation of membrane excitability.

**Figure 2.**
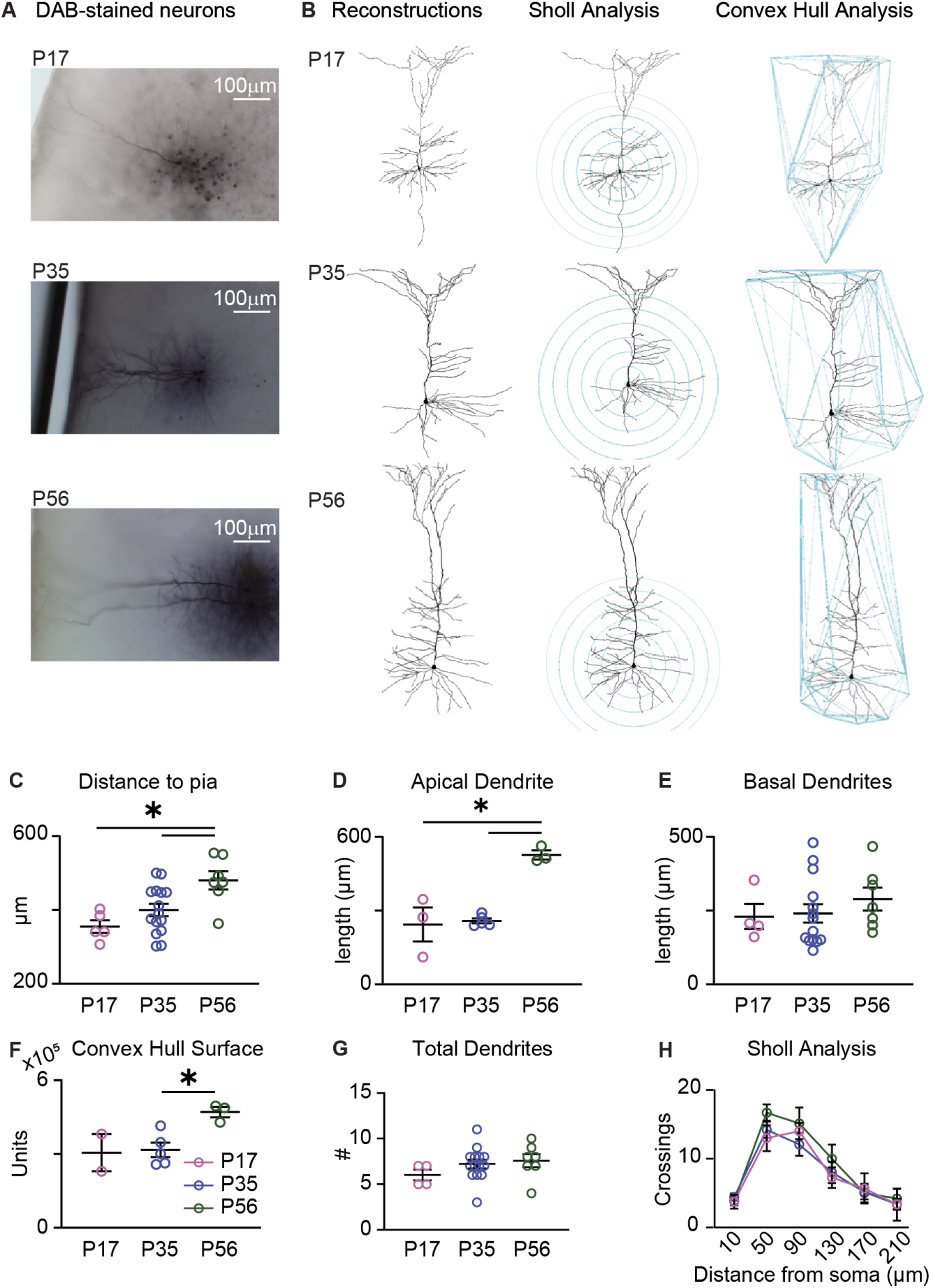
Increased apical dendrite and GC pyramidal neuron surface during postnatal development. **A.** Example DAB staining for L5 recorded neurons in different age groups. **B.** Neurolucida reconstructions of dendritic arbors, Sholl analysis and Convex Hull analysis for reconstructed neurons recorded from P17 (top), P35 (middle) and P56 (bottom). **C-E.** Plots of distance of soma from the pial surface (C), length of apical dendrite (D), and length of basal dendrites (E). **F.** Convex hull surface analysis for each age group. **G.** Total number of dendrites for each age group. **H.** Sholl analysis on reconstructed neurons by age groups. The data points represent average values by age group ± SEM. For C-G: black horizontal lines: average values ± SEM; open circles: single neuron measurements. For all panels: Pink: P17; blue: P35; green: P56. Asterisks: p ≤ 0.05.

### Developmental decrease in E/I balance onto GC pyramidal neurons

Previous work from other sensory cortices reported significant changes in synaptic transmission onto pyramidal neurons during postnatal development. While a previous study observed changes in synaptic transmission following sensory experience early in life in GC (Schiff et al., 2023), the information about developmental changes in synaptic drive onto GC neurons is lacking. To fill this gap in knowledge, we obtained voltage clamp recordings from GC infragranular pyramidal neurons in acute slice preparations obtained from mice at ages P17, P35, and P56. The morphology of recorded neurons was confirmed *post hoc*, as was the lack of expression of the inhibitory marker GAD67 (**Fig. 3A**). Spontaneous excitatory and inhibitory postsynaptic currents (sEPSCs and sIPSCs) were recorded by holding neurons at the reversal potential for chloride (-50mV in our experimental conditions, **Fig 3B**) and at the reversal potential for AMPA and NMDA receptor-mediated currents (+10mV in our experimental conditions, **Fig 3C**) respectively.

**Figure 3.**
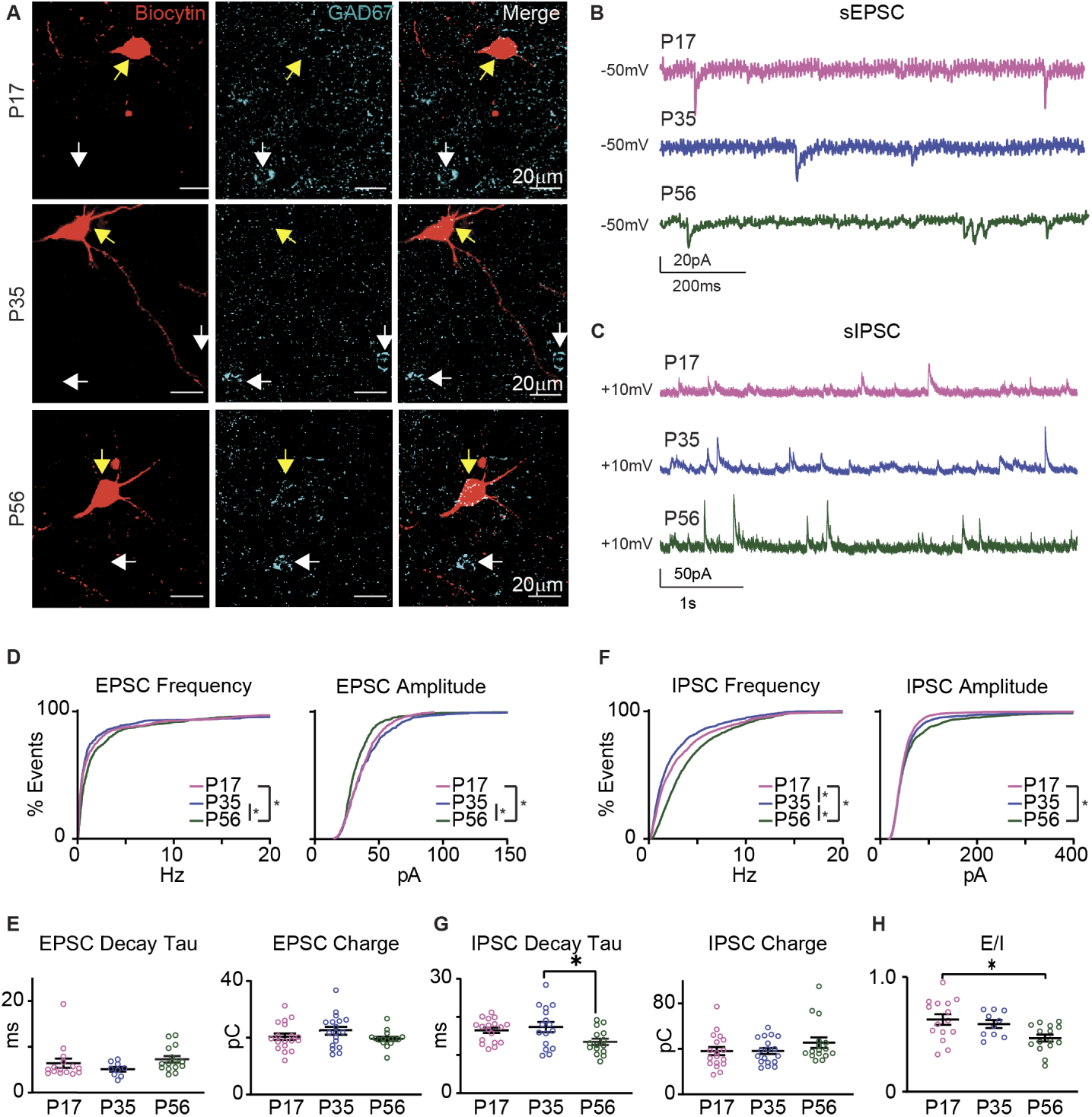
Shift in E/I balance during postnatal development. **A.** Sample single confocal plane images of biocytin-filled recorded neurons showing lack of colocalization with GAD67. The rows show a sample recorded neuron for each age. The yellow arrow points to the recorded neuron filled with biocytin (red); the white arrows point to examples of GAD67^+^ neurons at the same tissue depth. The lack of overlap indicates that the recorded neuron is GAD67^-^. **B.** Sample traces of sEPSCs recorded from P17 (pink), P35 (blue) and P56 (green) mice. **C. D.** Cumulative distribution of sEPSC frequency (left) and amplitude (right) by age group. P17: pink; P35: blue; P56: green. **E.** Plots of sEPSC decay time constant (tau) and sEPSC charge transfer. **F.** Cumulative distributions of sIPSC frequency (left) and amplitude (right) by age group. **G.** Plots of sIPSC decay time constant (tau) and sIPSC charge transfer. **H.** Ratio of sEPSC and sIPSC charge transfer for each age group. For all panels, black horizontal lines: average values ± SEM; open circles: single neuron measurements; asterisks: p ≤ 0.05.

Analysis of sEPSCs revealed opposite age-dependent changes in frequency and amplitude (**Fig. 3D**; P17 n=15 cells from 7 mice; P35 n=10 cells from 5 mice; P56=15 cells from 10 mice; **Fig. 3D left**, frequency: H=32.556 p=8.5e^-8^; P17 vs P35 p>0.999; P17 vs P56 p=5.566e^-6^; P35 vs P56 p=2.405e^-6^; **Fig. 3D right**, amplitude: H=35.272, p=2.191e^-9^; p17 vs p35 p>0.999; P17 vs P56 p=1.960e^-6^; P35 vs P56 p=8.860e^-7^; Kruskal-Wallis test with Dunn’s *post hoc* test). We did not observe a significant difference in sEPSCs decay time constant (tau; **Fig. 3E left**; P17 n=14 cells from 7 mice; P35 n=9 cells from 4 mice; P56 n=15 cells from 10 mice; F_(2,35)_=1.373 p=0.2667 one-way ANOVA) and no significant difference in the charge transfer for sEPSCs (**Fig. 3E right**; P17 n=18 cells from 8 mice P35 n=18 cells from 9 mice P56 n=16 cells from 8 mice, F_(2,49)_=1.604 p=0.212 one-way ANOVA), suggesting that the increase in sEPSCs frequency was likely compensated by the decrease in sEPSCs amplitude.

When assessing inhibitory synaptic drive, we observed that the frequency of sIPSCs was transiently reduced from P17 to P35 and then by P56 it increased to a level significantly higher than both P17 and P35 (**Fig. 3F**; P17 n=18 cells from 8 mice P35 n=18 cells from 9 mice P56 n=16 cells from 8 mice; **Fig. 3F left**, frequency: H=320.718 p<1.0e^-14^; P17 vs P35 p=2.907e^-6^; P17 vs P56 p<1.0e^-14^; P35 vs P56 p<1.0e^-14^ Kruskal-Wallis test with Dunn’s *post hoc* tests), suggesting a post-weaning transient decrease in inhibitory drive followed by an increase in inhibition by young adulthood. The amplitude of sIPSCs on the other hand showed a significant increase when comparing the youngest age group (P17) to P56 (**Fig. 3F right**, amplitude: H=14.23 p=8.0e^-4^; P17 vs P35 p=0.136; P17 vs P56 p=5.0e^-4^; P35 vs P56 p=0.205 Kruskal-Wallis test with Dunn’s *post hoc* tests), suggesting a gradual increase from before weaning to young adulthood. Analysis of decay kinetics showed a significant reduction in decay time constant (tau) selectively between P35 and P56 (**Fig. 3G left**; P17 n=18 cells from 8 mice; P35 n=16 cells from 9 mice; P56 n=15 cells from 8 mice; F_(2,46)_=4.000 p=0.0250 one-way ANOVA with Tukey’s *post hoc* tests: P17 vs P35 p=0.8054 P17 vs P56 p=0.0901 P35 vs P56 p=0.0259). This effect is consistent with previous reports from other brain regions which have associated with a shift in expression of GABA_A_ receptor subunits on pyramidal neurons (Heinen et al., 2004). The charge transfer of inhibitory synaptic events showed a trend toward increasing from P17 to adulthood; however, it was not statistically significant across age groups (**Fig. 3G right**; P17 n=18 cells from 8 mice P35 n=18 cells from 9 mice P56 n=16 cells from 8 mice F_(2,49)_=1.299 p=0.2821 one-way ANOVA). Overall, both frequency and amplitude of the sIPSCs showed a net increase from pre-weaning to young adulthood.

While on average the excitatory and inhibitory charge transfer was not significantly different, it showed opposite trends, suggesting that the E/I ratio may shift in the developmental window we investigated. We tested this possibility by calculating the ratio of excitatory and inhibitory charge (E/I ratio) for each recorded neuron and compared it across age groups. The E/I ratio was below 1 in all age groups, indicating that the balance between E/I balance favors inhibition in GC. While this parameter was stable between P17 and P35, it decreased significantly by P56 (**Fig. 3H**; P17 n=15 cells from 7 mice; P35 n=10 cells from 5 mice; P56 n=15 cells from 8 mice; F_(2,37)_=5.156 p=0.011 one-way ANOVA with Tukey’s *post hoc* tests: P17 vs P35 p=0.779; P17 vs P56 p=0.0098; P35 vs P56 p=0.1024), indicating that inhibition becomes even more dominant as neurons reach their mature state.

### Increased PV expression and accumulation of perineuronal nets during postnatal development

The increase in sIPSCs frequency and amplitude as well as the developmental shift of the E/I ratio favoring inhibition suggest a protracted maturation of inhibitory circuits in the deep layers of GC. Comparable changes have been observed in other cortical regions where they have been associated with PV^+^ INs (Heinen et al., 2004), a neuron type known to undergo several developmental changes in the temporal window we assessed in our experiments (Fagiolini and Hensch, 2000; Chattopadhyaya et al., 2004; Chattopadhyaya et al., 2007; Wu et al., 2012; Tatti et al., 2017). PV^+^ INs make synapses in the perisomatic region of pyramidal neurons (Chattopadhyaya et al., 2004; Kubota et al., 2016), making them the likely source of the majority of sIPSCs we recorded with patch clamp at the soma. Given these premises, we focused our analysis on this population of inhibitory neurons and asked whether known hallmarks of PV^+^ INs maturation could be detected in GC in the developmental window under study. Two well characterized markers of PV^+^ INs maturation are the expression of PV and the accumulation of perineuronal nets (PNNs) around PV^+^ INs somata (Pizzorusso et al., 2002; Lupori et al., 2023). We used immunohistochemistry with antibody staining against PV and the histological stain Wisteria Floribunda Agglutinin (WFA) to label PNNs in GC (**Fig. 4A**). We quantified the proportion of GC neurons that expressed PV, the overall proportion of PNNs and the proportion of PNNs on PV^+^ INs at P17, P35 and P56 using immunohistochemistry. The proportion of overall PV^+^ INs increased from P17 to P35, then remained stable to P56 (**Fig. 4B left**; P17 n=5 mice; P35 n=5 mice; P56 n=4 mice; F_(2,11)_=7.117 p=0.0104 one-way ANOVA with Dunnett’s *post hoc* tests: P17 vs P35 p=0.007; P17 vs P56 p=0.050). The overall proportion of PNNs in GC did not change across age groups (**Fig. 4B middle**, P17 n=5 mice; P35 n=5 mice; P56 n=4 mice; F_(2,11)_=1.077 p=0.3740 one-way ANOVA). However, the proportion of PNNs on PV^+^ INs increased significantly (**Fig. 4B right**: P17 n=5 mice; P35 n=5 mice; P56 n=4 mice; F_(2,11)_=7.597 p=0.0085 one-way ANOVA with Dunnett’s *post hoc* tests: P17 vs P35 p=0.0096; P17 vs P56 p=0.0158). These results suggest that both PV expression and association of PNNs with PV^+^ INs increase during a protracted period of postnatal development.

**Figure 4.**
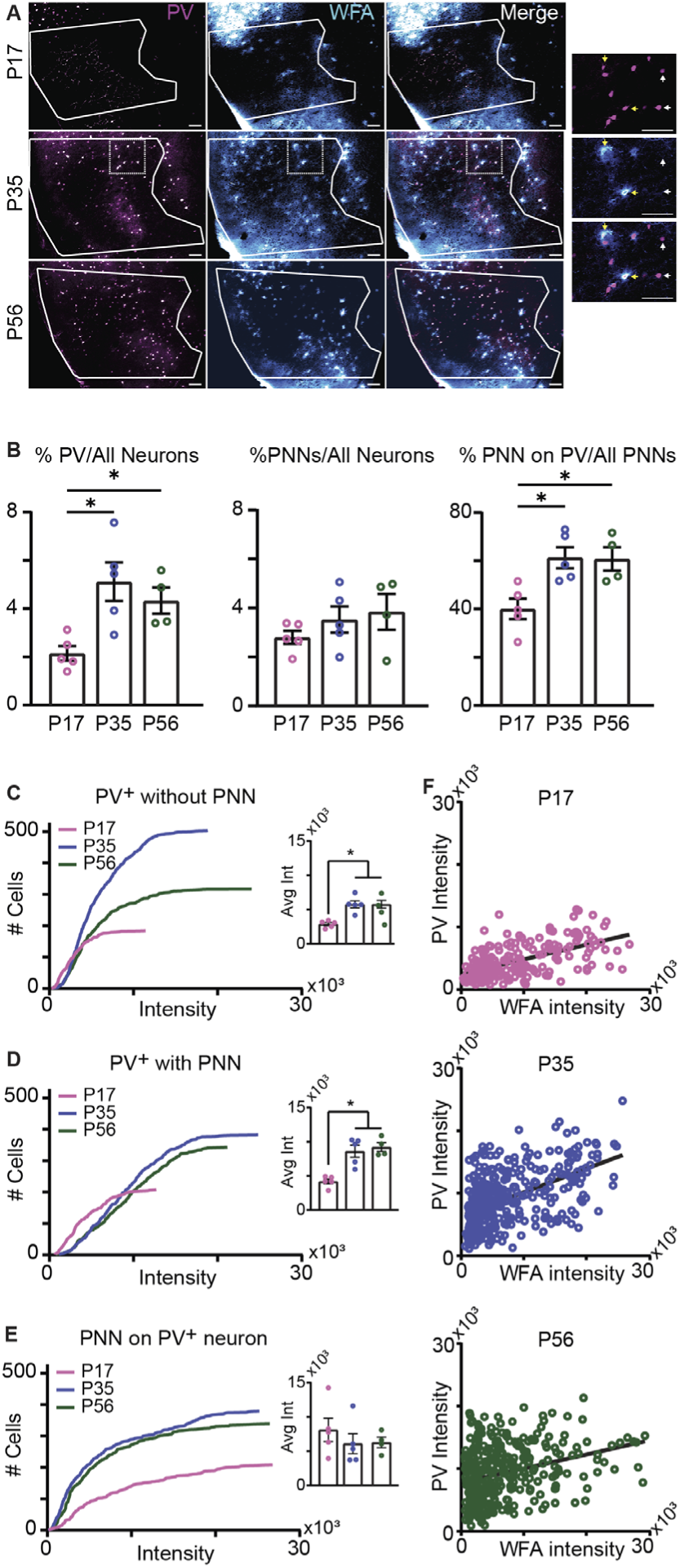
Increased PV expression and accumulation of PNNs during postnatal development. **A.** Sample images of GC slices processed for immunohistochemistry to label PV (magenta) and WFA to label PNNs (cyan). The region of interest containing GC is outlined in white. Sample images are comprised of z-stacks with 2µm step size taken at 10x magnification (scale bar: 100µm). The rows show PV, WFA, and a merged view for each age. The final column shows a zoomed in image of the boxed region outlined in the P35 sample. White arrows point to sample PV^+^ INs without a PNN; yellow arrows point to sample PV^+^ INs with a PNN. **B.** Left. Percentage of PV^+^ INs in GC in different age groups. Middle. Percentage of GC neurons with a PNN in different age groups. Right. Percentage of PNNs associated with PV^+^ INs. Open circles report counts by animal. Pink: P17; blue: P35; green: P56. **C.** Distribution of the intensity of the PV signal for PV^+^ INs without a PNN by age group. Note that the number of neurons and the intensity of PV show a transient increase at P35 before stabilizing at P56. The inset shows the average PV signal. **D.** Distribution of the intensity of the PV signal for PV^+^ INs with a PNN. Note that the signal reaches a steady state at P35. The inset shows the average signal. **E.** Distribution of the intensity of the WFA signal for PNNs associated with PV^+^ INs. Following an increase in fluorescence between P17 and P35 the signal stabilizes. The inset shows the average fluorescence signal. For C-E, open circles indicate counts for each animal. **F.** Correlation of PV intensity and WFA intensity for PV^+^ INs with a PNN. Open circles indicate single neuron signals. The solid line indicates a linear regression fit. Pink: P17; blue: P35; green: P56. Asterisks: p ≤ 0.05.

Next, we examined the intensity of the fluorescent PV signal in PV^+^ INs with and without a PNN during postnatal development and the fluorescence intensity of PNNs labeling on PV^+^ INs. The intensity of fluorescence of PV is associated with activity-dependent changes in expression (Donato et al., 2013) and the intensity of the WFA fluorescent signal informs on the degree of accumulation of a PNN (Slaker et al., 2016; Sorg et al., 2016; Lupori et al., 2023; Santos-Silva et al., 2024). For all age groups, we sorted PV^+^ INs into 2 groups, one without an associated PNN and one with a PNN. The fluorescence signal was quantified both as distributions vs cell number and as average signal (**Fig. 4C-E** and corresponding insets). In PV^+^ INs without a PNN, the distribution of PV signal intensity showed a progressive increase in PV fluorescence across age groups (**Fig. 4C**; PV^+^ INs without PNN: P17 n=184 cells from 5 mice; P35 n=503 cells from 5 mice; P56 n=317 cells from 4 mice; H=125.3 p<1.0e^-10^; P17 vs P35 p<1.0e^-10^; P17 vs P56 p<1.0e^-10^; P35 vs P56 p=0.3225; Kruskal-Wallis with Dunn’s *post hoc* test). The average intensity of the PV signal significantly increased from P17 to P35 (**Fig. 4C inset**; P17 n=5 mice; P35 n=5 mice; P56 n=4 mice; PV^+^ INs without PNN, F_(2,11)_=13.92 p=0.0010; P17 vs P35 p=0.0019; P17 vs P56 p=0.0028; P35 vs P56 p>0.999; one-way ANOVA with Tukey’s *post hoc* test) but there was no average change from P35 to P56, suggesting that the difference in signal intensity distributions from P35 to P56 is accounted for by an increase in the number of neurons expressing PV and not by an increase in average PV intensity.

Analysis of the same parameters for PV^+^ INs with a PNN showed an increase in PV intensity between P17 and P35 and no change between P35 and P56 (**Fig. 4 D**; PV^+^ with PNN: P17 n=208 cells from 5 mice; P35 n=382 cells from 5 mice; P56 n=343 cells from 4 mice; H=192.2 p<1.0e^-10^; P17 vs P35 p<1.0e^-10^; P17 vs P56 p<1.0e^-10^; P35 vs P56 p>0.9999; Kruskal-Wallis with Dunn’s *post hoc* test). Similarly, the average fluorescence signal significantly increased from P17 to P35 but showed no change from P35 to P56 (**Fig. 4D inset**; P17 n=5 mice; P35 n=5 mice; P56 n=4 mice; PV^+^ with PNN F_(2,11)_=1.008 p=0.0005; P17 vs P35 p=0.0015; P17 vs P56 p=0.0009; P35 vs P56 p=0.8053, one-way ANOVA with Tukey’s *post hoc* test), suggesting that after P35 the number of PV^+^ INs expressing PV has reached adult levels and that once they become associated with a PNN their PV expression is stable.

Finally, we quantified the fluorescence intensity of WFA labeling of PNNs associated with PV^+^ INs as a measure of PNN accumulation specifically on PV^+^ INs. There was an increase in the cumulative distribution of WFA fluorescence intensity after weaning, suggesting age-dependent aggregation of PNNs (**Fig. 4E**: P17 n=5 mice; P35 n=5 mice; P56 n=4 mice; H=21.35 p=0.2.314e^-5^; P17 vs P35 p=3.157e^5^; P17 vs P56 p=3.264e^-4^; P35 vs P56 p>0.999; Kruskal-Wallis with Dunn’s *post hoc* test). However, the average intensity of the WFA signal was not significantly different (**Fig. 4E**, inset: F_(2,11)_=0.4371 p=0.614 one-way ANOVA), suggesting that the developmental increase in WFA fluorescence depends on an increase in the number of PV^+^ INs associated with a PNN.

Finally, we asked whether the intensity of PV and WFA signals co-varied for each dual-labeled neuron within each age group (**Fig. 4F**). The PV and WFA fluorescence signals were significantly correlated in each age group (P17 r=0.556, p<0.001; P35 r=0.514, p<0.001; P56 r=0.307, p<0.001, Spearman correlation). The correlation did not increase with age, but the PV intensity signal increased, confirming the interpretation that PV expression increases after weaning and PNNs accumulate on PV^+^ INs as they reach their mature state by P56.

### Refinement of PV^+^ INs - pyramidal neurons connectivity during postnatal development

The developmental changes in inhibitory synaptic transmission we report in **Fig. 3** are likely dependent on changes in perisomatic inhibition, as sIPSCs originating from synapses around the soma of pyramidal neurons are better represented in patch clamp recordings. The increase in sIPSC frequency together with increase in PV immunofluorescence and increase accumulation of PNNs around PV^+^ INs suggest the possibility that the connectivity of PV^+^ INs onto pyramidal neurons undergoes refinement from pre-weaning to young adulthood. To assess possible developmental changes in connectivity, we took advantage of channelrhodopsin assisted circuit mapping (CRACM (Petreanu et al., 2007)). We injected a soma-restricted channelrhodopsin (soCoChr) into GC PV^+^ INs using the cre-lox system and stereotaxic surgery (Flex-soCoChr AAV injected into the left GC of PV-cre;Ai14 mice; **Fig. 5A**). One week after surgery we obtained coronal slices containing GC and performed patch clamp recordings from infragranular pyramidal neurons (**Fig. 5B**). After establishing the voltage clamp configuration, we used patterned light stimulation with a digital micromirror device (Polygon, Mightex) to shine blue LED light (473nm) onto spatially restricted regions of interest (30 ROIs approx. 100×100µm each) surrounding recorded neurons (approximately 500µm dorsal to ventral and 700µm from the bottom of layer 1 to the white matter/corpus collosum). We then determined the light-evoked response (or lack thereof) for each ROI (**Fig 5C**). As the number of PV^+^ INs at P17 was too low to effectively stimulate them with this approach, we compared local connectivity between P21 (viral injection at P14) and P56 (viral injection at P49). The location and morphology of recorded neurons was confirmed *post hoc*, and they were determined to be putatively pyramidal by their morphology, action potential properties, and absence of colocalization of biocytin with the tdTomato signal.

**Figure 5.**
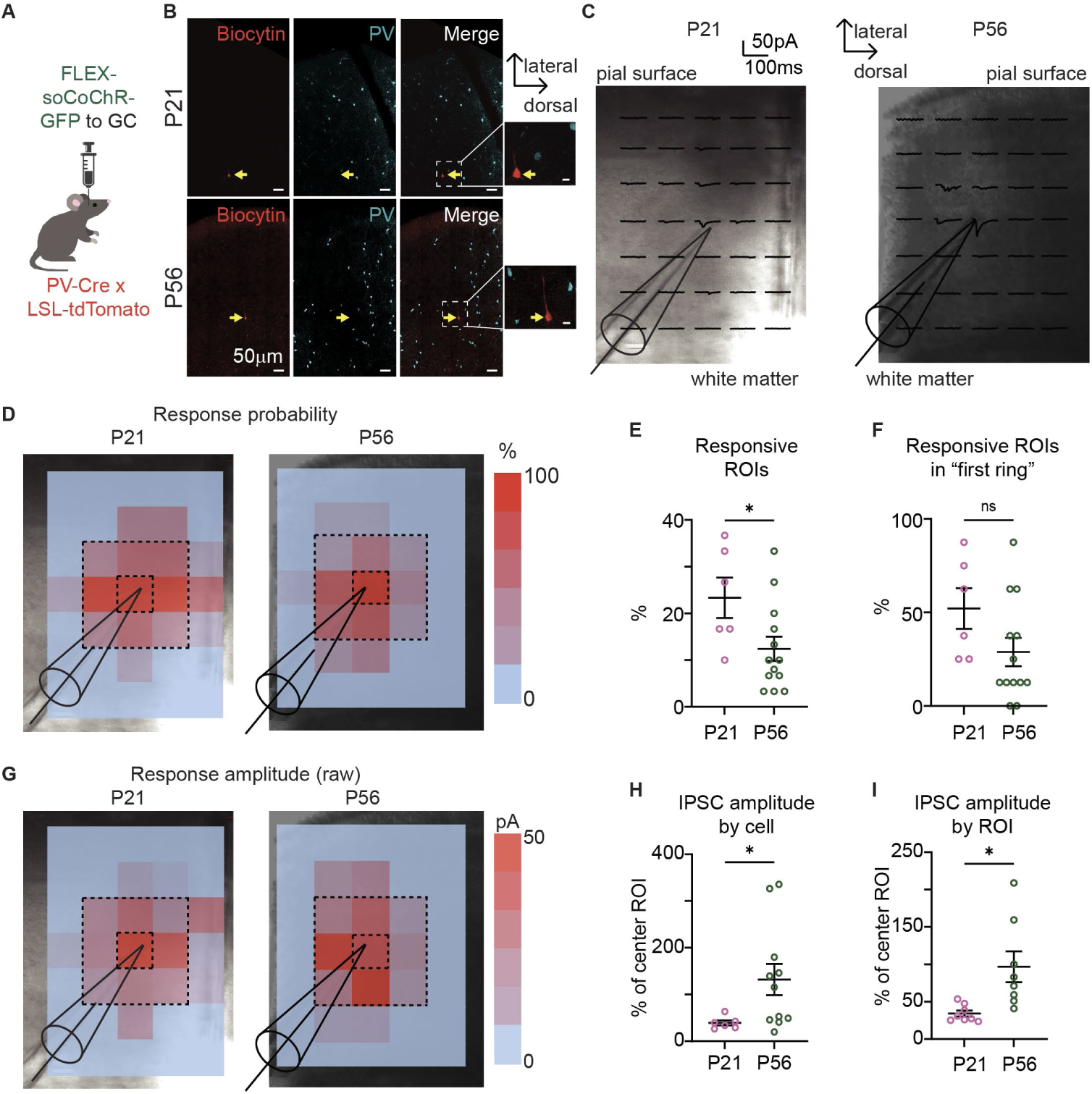
Refinement of PV connectivity and IPSC amplitude during postnatal development. **A.** Diagram of experimental approach. **B.** Sample images showing recorded neuron and ChR2-expressing PV neurons. Red: biocytin; cyan: ChR2-PV. Scale bar: 50 µm. Insets: zoomed in images of biocytin filled neurons. Scale bar: 10µm. **C.** Sample responses in 100µm x 100µm ROIs around a recorded neuron at P21 and P56. Tip of the recording electrode cartoon: position of recorded neuron’s soma. **D.** Sample heat map of the probability of evoking a response in each ROI. Tip of the recording electrode cartoon: position of recorded neuron’s soma. **E.** Plot of the percentage of stimulated ROIs (30 ROIs total) which evoked an IPSC. Open circles: values for each recorded neuron; black line: average value. **F.** Plot of the percentage of stimulated ROIs within the “first ring” (8 ROIs total), which evoked an IPSC. Open circles: values for each recorded neuron; black line: average value. **G.** Sample heat map of IPSC amplitude in each ROI. **H.** Plot of IPSC amplitude for stimulated ROIs in the “first ring” presented as % of the amplitude of the IPSC evoked by stimulating the ROI containing the soma of the recorded neuron. Normalized IPSC amplitudes are presented as average for each cell. Open circles: values for each recorded neuron. **I.** Amplitude of IPSCs by age group, represented as % of the IPSC evoked by stimulation of the ROI containing the soma of the recorded neuron. Data are presented as average for each ROIs (8 ROIs total). Open circles: values for each ROI. For all panels: pink: P21; green: P56; black horizontal lines: average values ± SEM; open circles: single neuron (or ROI) measurements; asterisks: p ≤ 0.05.

To be included in the analysis, neurons had to respond with an IPSC to light stimulation in at least 1 ROI (**Fig. 5D**). For each recorded neuron, we first analyzed the percentage of ROIs that, when stimulated, evoked a detectable IPSC (out of 30 ROIs/pyramidal neuron). Heatmaps show percent success in evoking an IPSC in each ROI quantified across all neurons by age group. We found that the number of ROIs that when stimulated evoked IPSCs in neurons from P56 mice was smaller compared to P21 (**Fig 5E** P21: n=6 cells from 3 mice, IPSCs evoked in 7.0±1.29 out of 30 ROIs; P56: n=13 cells from 6 mice, IPSCs evoked in 3.69±0.91 out of 30 ROIs; U=15 p=0.0331 Mann-Whitney U test). We then restricted the analysis to the ring of ROIs immediately surrounding the soma of the recorded neuron and compared responsiveness in this “first ring” of ROIs (8 total ROIs, indicated by the dashed line in Fig 5D). We observed a trend toward reduced connectivity at P56 that did not reach significance (**Fig 5F**; P17: 4.17±0.87 out of 8 ROIs, P56: 2.3±0.9 out of 8 ROIs; U=18.5 p=0.0659 Mann-Whitney U test). These findings indicate that the connectivity of GC PV^+^ INs clusters around the perisomatic region of infragranular pyramidal neurons and suggest that PV^+^ INs input onto GC pyramidal neurons becomes more spatially restricted over the course of postnatal development.

The spatial restriction of connectivity around the perisomatic area at P56 correlates with the postnatal time window showing an increase in sIPSCs frequency and amplitude (see **Fig. 3**). We, therefore, tested the possibility that this refinement in connectivity may be accompanied by an increase in amplitude of evoked IPSCs. To account for differences in virus expression and light stimulation intensity across preparations, the amplitude of evoked IPSCs by stimulation of each ROI was normalized to the amplitude of the IPSC evoked by stimulating the ROI containing the soma of the recorded pyramidal neuron. This analysis was restricted to responsive ROIs in the “first ring” indicated by dashed lines in Fig 5G, as they showed the highest response probability in both age groups. Heat maps of IPSCs amplitudes (**Fig. 5G**) show differences in the raw IPSC amplitude by ROIs. To compare IPSC amplitudes across neurons we normalized data in two ways. In Figure 5H we report IPSC amplitude normalized to the IPSC evoked in the center ROI for each recorded neuron. IPSCs recorded in P56 mice were significantly larger compared to P21 mice (**Fig. 5H**; P21 n=6 cells from 3 mice; P56 n=13 cells from 6 mice; U=11 p=0.0273 Mann-Whitney U test). While at P21 the IPSC in the central ROI was larger than the IPSCs in the surrounding ring of ROIs, at P56 this was not always the case and with IPSCs larger in the “first ring” of ROIs compared to the central ROI. To explore this possibility further, we plotted the normalized IPSCs by ROI to test for potential confounds due to the location of the stimulation (**Fig. 5I**). Again, at P56 we observed more ROIs with IPSCs larger than the IPSC recorded in the central ROI (n=8 ROIs for each age group; U=3 p=0.0011 Mann-Whiteny U test), confirming the age-dependent increase in IPSC amplitude in the perisomatic region of pyramidal neurons. Together, these results indicate that during postnatal development the connectivity of GC PV^+^ INs onto infragranular pyramidal neurons is refined to become spatially restricted around the soma, while inhibitory synaptic drive increases. This effect is consistent with the increase in inhibitory drive shown in Fig. 3 and with a progressive decrease in excitability of pyramidal neurons over the course of postnatal development.

## Discussion

Our results show that the developmental window from P17 to P56, a sensitive window for the development of taste preference (Schiff et al., 2023), is characterized by a developmental decrease in the excitability of pyramidal neurons in the infragranular layers of GC. This effect is associated with subtle changes in intrinsic membrane properties, with a shift of the E/I balance of overall synaptic drive toward inhibition and a refinement in connectivity and synaptic strength of inhibitory inputs mediated by PV^+^ INs. The temporary decrease in intrinsic excitability of pyramidal neurons in the fifth postnatal week was characterized by a change in the input/output function between P17 and P35. Similar effects were recently reported in mouse primary visual cortex (V1) at the transition from eye opening to the peak of the critical period for ocular dominance plasticity (Ciganok-Huckels et al., 2023), suggesting that this may be a common feature of sensory circuits. In V1, the shift in input/output function is permanent (Ciganok-Huckels et al., 2023), while in GC, the decrease we observed was transient, as the input/output function of pyramidal neurons at P56 was comparable to that of P17 mice. Another important difference between developmental changes in pyramidal neurons between GC and other cortical areas is that the decrease in excitability was not accompanied by changes in rheobase and action potential threshold, as observed in V1 (Ciganok-Huckels et al., 2023) and prefrontal cortex (Kroon et al., 2019). Interestingly, in contrast to the recovery of input/output function by P56, GC infragranular pyramidal neurons showed a persistent decrease in input resistance between P17 and P56. This change in input resistance may be accounted for by an increase in the complexity or size of the pyramidal neuron, as we observed an increase in the apical dendrite length and convex hull analysis. An increase in pyramidal neurons size was previously observed in the prefrontal cortex, although it occurred much earlier in development (Kroon et al., 2019). In addition, our findings in GC also differ from the morphological changes reported for V1, where basal dendrites, but not apical dendrites, increased in size (Ciganok-Huckels et al., 2023), suggesting region-specific changes in the laminar distribution of inputs onto pyramidal neurons. Taken together, our results are in line with findings from other cortical regions in reporting an overall decrease in deep layer pyramidal neurons intrinsic excitability and changes in pyramidal neurons morphology between juvenile and young adult mice during postnatal development and suggest the engagement of region-specific mechanisms.

In addition to the changes in intrinsic excitability, we observed a shift in the E/I balance of synaptic charge that was associated with an opposite shift in sEPSCs amplitude and frequency and with an increase in both sIPSCs amplitude and frequency. The changes in synaptic drive appear delayed compared to the shift in input/output function. Indeed, sEPSCs and sIPSCs amplitude and frequency were comparable between P17 and P35, while both parameters were significantly different at P56, suggesting that the postnatal development of synaptic function in GC extends beyond the 5^th^ postnatal week. The developmental decrease in sEPSC amplitude was accompanied by an increase in sEPSC frequency. While this effect is comparable to previous reports from V1 (Desai et al., 2002; Tatti et al., 2017), the changes in GC occur over a more extended developmental window. The sEPSCs decay kinetics were stable, as was the total excitatory charge transfer, suggesting that the excitatory drive onto infragranular GC pyramidal neurons is overall stable between pre-weaning and adulthood.

Differently, analysis of sIPSCs revealed an increase in both frequency and amplitude, accompanied by a decrease in the decay kinetics from pre-weaning to adulthood. The faster sIPSC decay kinetics may partially compensate for the increase in amplitude and frequency, as the overall charge transfer for inhibitory events was not significantly different during the developmental window under study. Nonetheless, the increase in frequency and amplitude of sIPSCs in the face of overall stable excitatory drive resulted in a significant shift in the E/I balance of charge transfer towards inhibition. This effect, together with the decrease in input resistance, renders GC infragranular neurons less excitable in adulthood compared to the pre-weaning period and points to a powerful inhibitory control of GC output.

In rodent V1, a decrease in kinetics in the presence of increased amplitude is associated with a switch in prevalence of GABA_A_ receptors containing the α3 subunit in favor of GABA_A_ receptors containing α1. This switch in subunits in V1 is complete by the third postnatal week and is an important mechanism regulating the process of maturation of inhibitory circuits (Heinen et al., 2004). Different from V1, the acceleration of sIPSC decay kinetics in GC occurs between the fifth and seventh postnatal week, raising the possibility that the maturation of inhibitory synapses in this brain region extends into adulthood. In addition to changes in sIPSC amplitude and kinetics, we observed a significant increase in sIPSC frequency by P56, which is typically associated with modulation of presynaptic release (Zucker and Regehr, 2002).

In patch clamp recordings from pyramidal neurons somata, the majority of sIPSCs are generated by perisomatic inputs, which are primarily from PV^+^ INs. This population of neurons is well known to undergo extended postnatal refinement of connectivity and synaptic strength. The number of neurons expressing PV increases during the postnatal period (Gonchar et al., 2007; Tatti et al., 2017) and changes in PV expression is associated with learning (Donato et al., 2013) and experience-dependent plasticity in the gustatory cortex (Schiff et al., 2023).While the expression of PV increases after the second postnatal week, PV^+^ INs become associated with PNNs whose accumulation reaches stable levels around the closure of the sensitive windows for experience-dependent plasticity (Pizzorusso et al., 2002; Berardi et al., 2003; Pizzorusso et al., 2006; Sorg et al., 2016). In GC, our data show that the expression of PV and accumulation of PNNs on PV^+^ INs increased in a correlated fashion between pre- and post-weaning, similar to previous reports in other cortices (Lupori et al., 2023). Interestingly, in GC, the accumulation of PNNs around PV^+^ INs is complete before the increase in inhibitory synaptic drive, further supporting the interpretation that in this cortical region the maturation of synaptic transmission extends over a longer postnatal time window compared to other regions (Lo et al., 2017; Tatti et al., 2017)).

Analysis of the connectivity of PV^+^ INs onto infragranular pyramidal neurons highlights a process of refinement characterized by restriction of connectivity to the perisomatic region, suggesting post-weaning pruning of inhibitory synapses and increase in synaptic efficacy. Interestingly, this process of refinement follows a similar temporal window as the increase in sIPSCs frequency and amplitude we report in Fig. 3. Previous work in organotypic cultures of V1 identified a postnatal increase in GABA with increased pruning of PV^+^ INs synapses leading to a restriction of PV^+^ INs innervation around the perisomatic region of pyramidal neurons (Chattopadhyaya et al., 2007; Wu et al., 2012). As changes in frequency of spontaneous synaptic currents are associated with changes in neurotransmitter release (Zucker and Regehr, 2002), it is tempting to speculate that the developmental increase in sIPSCs frequency and amplitude depends on a combination of increase in GABA release and refinement of PV^+^ INs connectivity in the perisomatic region of GC infragranular pyramidal neurons.

The picture emerging from our analysis points to an extended postnatal process of maturation of cortical circuits for taste that straddles the developmental window between weaning and adulthood. This process involves changes in membrane properties and a shift in E/I balance toward inhibition of synaptic drive onto infragranular pyramidal neurons. In parallel, inhibitory circuits mediated by PV^+^ INs are refined to focus their influence on the perisomatic region of infragranular pyramidal neurons, possibly modulating the output of GC. The timeline of GC maturation overlaps with a developmental window during which experience with tastants has a lasting influence on taste preference that persists into adulthood (Schiff et al., 2023). Our findings suggest that the transition to independent feeding marks a highly plastic time in the development of taste processing that can profoundly influence eating habits in adults (Schiff et al., 2023). When viewed in the context of previous work on postnatal maturation of cortical circuits (Micheva and Beaulieu, 1995; Fagiolini and Hensch, 2000; Chattopadhyaya et al., 2004; Gogolla et al., 2014; Lo et al., 2017; Tatti et al., 2017; Takesian et al., 2018; Ciganok-Huckels et al., 2023; Schiff et al., 2023), our study highlights common features, some of which occur over comparable developmental windows, supporting the interpretation that these may be general mechanisms for neural circuit function. However, the events underlying the maturation of cortical circuits appear to occur over region-specific timelines, with GC circuits maturation extending over a longer postnatal window compared to somatosensory, auditory and visual cortical circuits (Micheva and Beaulieu, 1995; Lo et al., 2017; Tatti et al., 2017; Takesian et al., 2018; Ciganok-Huckels et al., 2023). This feature of GC circuits may depend on the need for extended experience-dependent plasticity for taste to allow for the acquisition of the identity of safe and nutritious food while foraging, a process that engages other sensory and motor systems to find nutrients.

## Author Contributions

Designed experiments: HS, AM; Performed experiments: HS; Analyzed data: HS Supervised experiments and analysis: AM; Writing: HS, AM

## Acknowledgments

We wish to thank Mark Bodik for help with the morphological reconstructions and Maria Isaac and Aylar Berenji-Kalkhoran for support with imaging.

## Data Availability

All data needed to evaluate the conclusions in the paper are included in the manuscript.

